# Environmental enrichment reverses memory impairment in B3-ARKO mice

**DOI:** 10.1101/2020.08.04.234849

**Authors:** Thais Terpins Ravache, Gabriela G. Nunes, Alice Batistuzzo, Fernanda B. Lorena, Bruna P. P. do Nascimento, Martha Bernardi, Miriam O. Ribeiro

## Abstract

Norepinephrine plays an important role in modulating the processes of memory consolidation and evocation through its beta-adrenergic receptors (Adrβ), which are expressed in the hippocampus and amygdala. Several studies have shown that all three subtypes of Adrβ (β_1_, β_2_ and β_3_) play an important role in cognition. Environmental enrichment (EE), a technique initially used to decrease the stress of animals held in captive environments, has also been shown to produce cognitive benefits in both healthy and sick animals. In this study, we hypothesized that EE would reverse the memory impairment induced by the absence or Adrβ_3_. To test this, 21- and 86-day-old Adrβ_3_KO mice were exposed to an EE protocol for 8 weeks. The study showed that the EE protocol is able to correct the memory impairment when applied to Adrβ_3_KO animals immediately after weaning but has no effect when applied to adult animals. We also found that aging worsens the memory of Adrβ_3_KO mice. Our results suggest that a richer and more diverse environment helps to correct memory impairment in Adrβ_3_KO animals. They also reinforce the idea that noradrenergic signaling is involved in the cognitive impairment observed late in life, as aging led to a worsening in the memory of the Adrβ_3_KO animals that was not corrected by the environmental enrichment protocol.

## INTRODUCTION

Norepinephrine (NE) plays an important role in modulating the processes of memory consolidation and evocation through its beta-adrenergic receptors (Adrβs), which are expressed in the hippocampus and amygdala of mammals(1, 2). Adrβ_1_ is found in the hippocampal regions CA1 and CA3 and in a small amount in the dentate gyrus. Adrβ_2_ and Adrβ_3_ are found in practically all regions of the hippocampus and amygdala, with β_3_-AR also being present in the entorhinal cortex (3–5). Adrβ selective agonists stimulate the induction of long-term potentiation (LTP) in the pyramidal cells of the CA1 region of the hippocampus and are involved in hippocampal-dependent cognitive functions (6). The use of Adrβ antagonists and the blockage of NE synthesis impair the formation of hippocampal-dependent memory (6).

Since Adrβs are positively coupled to protein G and enable the amplification of neuronal signals (2), Adrβs mediate the formation of long-term memory, probably by activating cAMP signaling pathways with consequent modulation of neuronal plasticity and excitability (7, 8).

Isopropoterenol, an agonist for Adrβ_1_ and Adrβ_2_, increases neural plasticity in the CA1 and CA3 regions and in the dentate gyrus in the hippocampus, while propranolol, a Adrβ_1_ and Adrβ_2_ antagonist, blocks LTP in the CA1 region and the dentate gyrus (9–12). In addition, the absence of the Adrβ_3_ receptor induces important deficits in the formation of short- and long-term memory (13).

Environmental enrichment (EE) is a technique that was initially used as a way of decreasing the stress of animals held in captive environments (14) (15). However, many studies about strategies for investigating cognitive skills have shown that EE has cognitive benefits in animals whether they are healthy, sick, young or old (15–17). EE may include 1) Physical enrichment, in which there is the introduction of materials in the enclosure that stimulate activities in the animal’s natural environment, such as burrows and tunnels; 2) Sensory enrichment, which consists of stimulating the animal’s senses with herbs to stimulate the sense of smell, and objects with different textures for tactile stimulation; 3) Cognitive enrichment, activities in which the animal needs to solve a problem, for example, hanging a banana on a string through the roof of the cage will require some increased cognitive demand to solve the challenge and reach the prize; 4) Social enrichment, animals can be allowed interact with other species, or the number of animals of the same species in the same enclosure can be increased which will consequently influence the social hierarchy; 5) Food enrichment, which consists of changes in the animals’ diet or changes in the frequency or time of feeding (14, 18).

Several studies evaluating the effect of EE on brain development have observed increased neurogenesis, increase in dendritic ramifications, as well as increased nerve growth factor (NGF) gene expression and increased LTP in the hippocampus (19–21). EE has also been shown to improve learning and memory consolidation in animal models of Alzheimer’s disease (22, 23).

Here we hypothesized that the use of EE could reverse the memory impairment induced by the absence of Adrβ_3_. Thus, the aim of our study was to evaluate the effect of EE on memory consolidation processes in Adrβ_3_ knock-out (Adrβ_3_KO) mice.

## METHODS

### Animals

Adrβ_3_KO mice with an FVB background, generated by removing the 306bp genomic fragment containing the sequences encoding the third through the fifth transmembrane domains of the Adrβ_3_ and replacing it with a neomycin selection cassette, as described by Susulic et al. (24), were obtained from Mackenzie Presbyterian University (Sao Paulo, Brazil). The animals were genotyped to confirm their status as homozygous knockout (β_3_-ARKO) or wild type (WT) mice. Male Adrβ_3_KO mice and WT controls from different litters randomized between groups were used in a protocol approved by the Institutional Committee on Animal Research at the Center of Biological Sciences and Health, Mackenzie Presbyterian University. Each experiment was repeated twice on the four different groups of animals. Mice were housed in groups at 26°C, 55–60% humidity, and a 12-h light/dark cycle with *ad libitum* access to standard food (Nuvilab, Brazil) and water.

### Experimental design

Based on studies on mice development, we evaluated the effects of EE at two moments in the lives of animals. Study 1 assessed the impact of EE that started right after weaning, PND21 until PND85. Study No. 2 assessed the impact of EE initiated in adult life, so EE started on PND120 until PND 180.

### Study 1 - Effect of early EE on young mice

The animals were transferred immediately after weaning on post-natal day 21 (PND21) to the EE cage and were submitted to the protocol described in Table 1 until PND85 when behavioral tests were started and finished at PDN120 (Figure 1A). The animals were divided into the following groups: WT (n = 7); Adrβ_3_KO mice (n = 7); WT + EE (n = 9); and Adrβ_3_KO + EE (n = 9).

**Figure 1.**
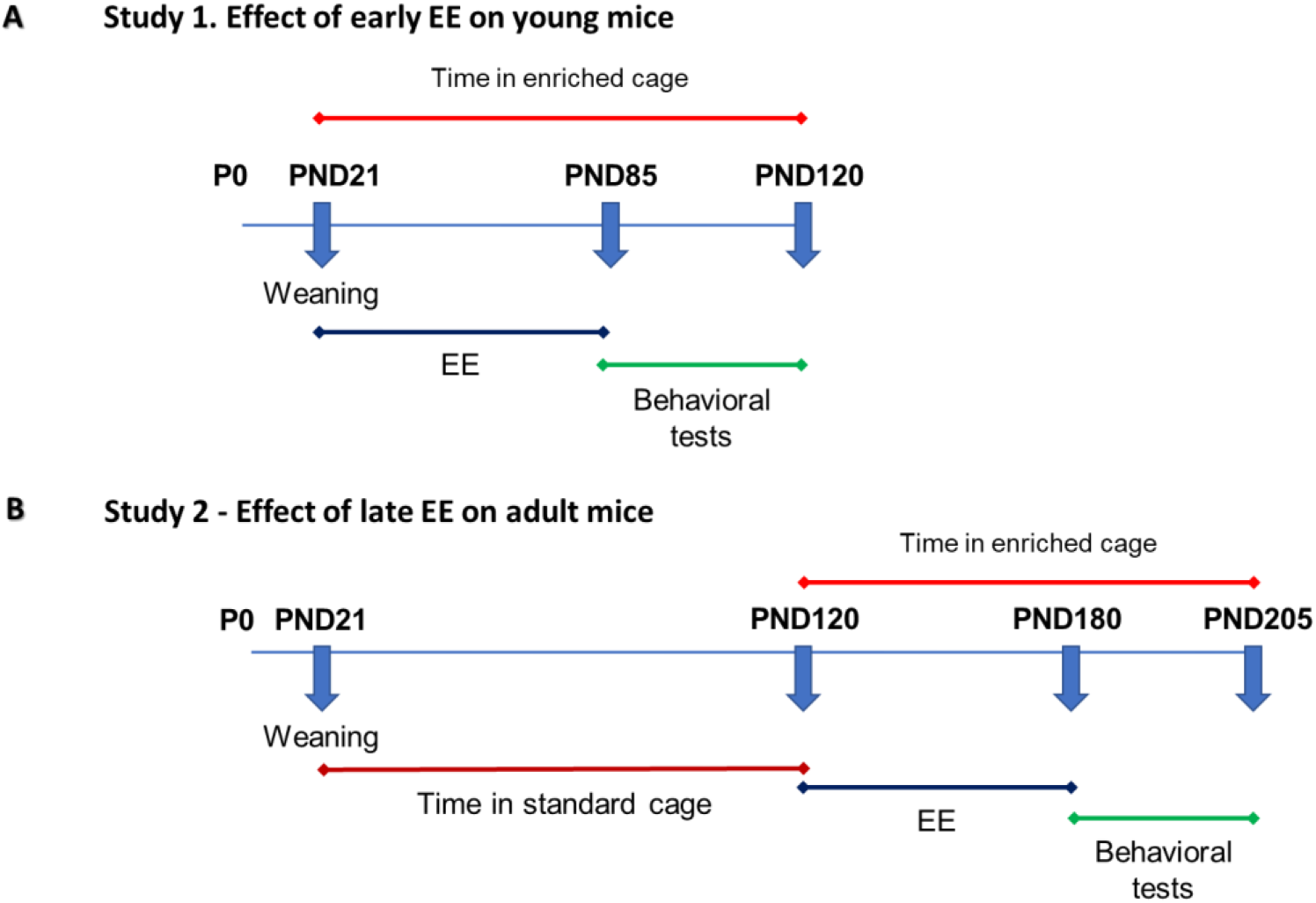
Experimental design. A) Study n°1: Effect of EE started right after weaning on post-natal day 21 (PND21) until PND85. The behavior assessment started on PND85 until PND120. During the tests the animals remained in the enriched cage but without the interventions B) Study n° 2: Effect of EE initiated in adult life on PND120 until PND 180 when the behavior assessment was performed until PND 205.

**Table 1:** EE protocol.

| Weeks | 1<sup>st</sup> Intervention | 2<sup>nd</sup> Intervention |
| --- | --- | --- |
| 1<sup>st</sup> | Familiarization with the new environment. | Fruit salad (banana, apple, grape) for 5-6 hours |
| 2<sup>nd</sup> | Exposure to cotton balls of different sizes for 4-5 hours | Hiding fruits under the bedding for 4-5 hours |
| 3<sup>rd</sup> | Exposure to ice with and without water for 1 hour | Exposure to newspaper sheets for 5-6 hours |
| 4<sup>th</sup> | Exposure to carrots in different sizes for 4-5 hours | Exposure to plastic balls in a box filled with bedding for 3-4 hours |
| 5<sup>th</sup> | Exposure to trail with seasonings (oregano, lemongrass and chamomile) for 2 hours | Exposure to cooked rice for 3-4 hours |
| 6<sup>th</sup> | Exposure to two extra burrows made from cardboard. | A banana hanging from a thread attached to the roof of the cage for 2-3 hours |
| 7<sup>th</sup> | Exposure to mirrors for 15 minutes | Exposure to neutral jelly with raisins inside for 3 hours |
| 8<sup>th</sup> | Exposure to bedding from a rat cage in placed in four different locations in the cage for 1 hour | Exposure to bowls containing water with frozen peas, carrots and corn for 2-3 hours |

### Study 2 - Effect of late EE on adult mice

The animals were kept in standard cages until PND120, when they were then transferred to the EE cage and submitted to the EE protocol until PND180 (Figure 1B). Behavioral tests were started at PND180 and finished at PDN 205. The animals in this study were divided as in Study 1: WT (n = 6); Adrβ_3_KO (n = 9); WT + EE (n = 7); and Adrβ_3_KO + EE (n = 7).

### EE Protocol

All the mice submitted to the EE remained in two-story cages (57×31×41cm), lined with wood shavings and with a shelter, water and food on both floors. The EE protocol was standardized in our laboratory (adapted from (25, 26)) and consisted of two interventions per week for eight weeks in the morning, with sensory, cognitive and dietary activities, as well as changes in the home cage (Table 1). After eight weeks of EE, the behavioral tests were started. During the behavioral assessment the animals remained in the EE cages until the completion of the tests, but without the stimulatory activities.

### Behavioral testing

All tests were performed in the morning (7:00–9:00 AM), under dimmed light (15 lux), and recorded by video for later analysis for two different blind observers in the following order for both studies 1 and 2:

### Open field test (OF)

The open field test was used to evaluate exploratory activity (27). The animals were placed in the center of a circular acrylic arena (diameter = 30cm) divided into four central zones and eight peripheral zones (Insight Ltda, Brazil), in a low-light environment (15 Lux) for 10 minutes. Locomotion (total number of lines crossed with all four paws) in the central and peripheral zones was measured. The test was performed three consecutive times with a 24-hour interval (28).

### Novel object recognition test (NOR)

This test was performed to evaluate short- and long-term memory. It was performed in the OF arena right after the OF test in order to guarantee the habituation of the mice to the arena. The test consists of three stages: familiarization to the two unknown objects, 3 and 24 hours after the familiarization. In the familiarization stage, the animals were placed in the open field arena for 10 minutes. After familiarization, the animals were exposed to two unknown and identical objects, object O1 and object O1’ for 3 minutes. Three hours later, the test was performed with the animals being placed in the arena for 3 minutes and exposed to object O1 and a new object (O2). 24 hours after the familiarization the animals were placed in the arena for 3 minutes and exposed to the known object O1 and a new object (O3). At each stage the time spent with objects, i.e. animal exploring the object with their nose, was expressed as a recognition index, i.e., the percentage of time spent with each object considering the total time spent with both objects (29).

### Social recognition test (SR)

Social preference and discrimination were evaluated using a non-automated 3-chambered box with three successive and identical chambers (Stoelting, Dublin). The protocol used is similar to the one described previously (30). Briefly, in the familiarization period, the mice were allowed to explore the three chambers freely for 10 min starting from the intermediate compartment, with the two other chambers containing empty wire cages. To test social preference, the test mouse was placed in the intermediate compartment, while an unfamiliar mouse was now put in one of the wire cages in a random and balanced manner. The doors were re-opened and the test mouse was allowed to explore the three chambers for 10 min. Time spent in each of the chambers, the number of entries into each chamber, and the time spent sniffing each wire cage were recorded for social preference. In the third phase, social discrimination was evaluated with a new stranger mouse (unknown) being placed into the remaining empty wire cage with the test mouse allowed to explore the entire arena for 10 min, having the choice between the first, already-investigated mouse (known) and the novel unfamiliar mouse (unknown). The same measures were taken as for the social preference (31) (32).

### Statistical analysis

Experimental data were analyzed using PRISM software (GraphPad Software). The statistical significance of the difference among the mean values for the groups were analyzed by two-way ANOVA, followed by the Tukey’s test was used, with a significance level of p ≤ 0.05.

## RESULTS

### EE exposure early in life increases ambulatory activity in young adult B3-ARKO and WT mice

In the OF the 2-way ANOVA showed that control Adrβ_3_KO young mice exhibited less ambulatory activity when compared to adult WT young mice (*F*(1,6) =12.63; p=0.012), but there was a significant reduction of ambulatory activity after the 3 days exposure in OF for both groups (*F*(2,12)=46.41; p<0.0001) (Figure 2A). It was noted that after the EE exposure the difference between WT and Adrβ_3_KO young mice in total line crossing disappears (*F*(2,32)=2.16; p=0.13) and both groups exhibited a reduction in ambulatory activity after the three days of exposure to the OF (*F*(1,22)=31.02; p<0.0001) (Figure 2B). Regarding the exploratory activity there was an effect of treatment only on the 3^rd^ day of exposure to the open field test (*F* (1,28) = 17.55; p=0.003) (Figure 2C-E).

**Figure 2.**
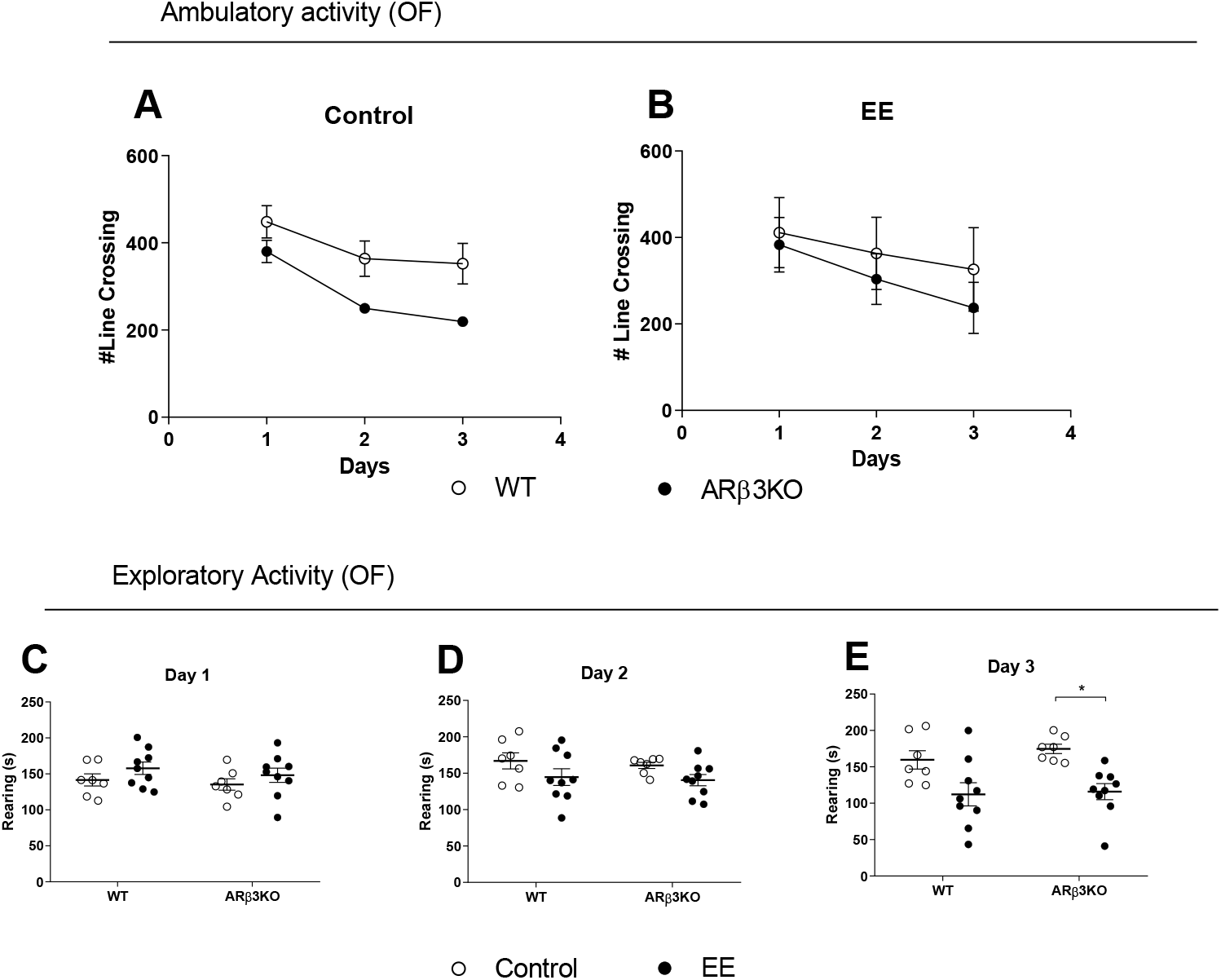
Effect of exposure to EE early in life on locomotor activity and anxiety behavior in young adult B3-ARKO and WT mice. *Open Field*. **(A)** Total number of line crossing of WT and ARB3KO without exposure to EE (WT vs. Adrβ_3_KO with p=0.01; Day 1 vs 2 and 3 with p<0.0001); **(B)** Total number of line crossing of WT and ARB3KO exposed to EE (WT vs. Adrβ_3_KO with p=0.13; Day 1 vs 2 and 3 with p<0.0001); **(C-E)** Total number of rearing for control WT vs. WT EE young mice on Day 1 (p=0.38; t=1.3), Day 2 (p=0.21; t=1.66) and Day 3 (p=0.03; t=2.64); Total number of rearing for control Adrβ_3_KO vs. Adrβ_3_KO EE young mice exposed to EE on Day 1 (p=0.65; t=0.99), Day 2 (p=0.28; t=1.51) and Day 3 (p=0.005; t=3.28).Values are expressed as mean ± SEM (n=10). * p< 0.05.

### EE exposure early in life corrects cognitive impairment in young adult B3-ARKO mice

Cognition was evaluated through the novel object recognition test (NOR) and the valence-based social recognition test (SR), which are based on exploratory behavior and assess memory and preference for novelty. In the NOR test, all groups explored the objects similarly during the familiarization period (Supp Fig. 1A-B). WT mice spent significantly more time with the new object, 3h (O2) (p<0.0001; t=5.06) and 24 h (O3) (p<0.0001; t=8.52) after the familiarization period (Figures 3 A-B). In contrast, the Adrβ_3_KO spent similar amounts of time with old (O1) and new objects (O2) 3 h (p=0.62; t=1.03) and 24h (O3) (p=0.1; t=2.06) after the familiarization period (Figure 3 A-B), confirming data from a previous study (13) showing that the absence of Adrβ_3_ impairs memory consolidation. Remarkably, EE was effective in correcting the memory impairment displayed by the Adrβ_3_KO mice both 3h (O2) (p<0.0001; t=8.06) and 24 h (O3) (p=0.0014; t=3.73) after the familiarization period (Figure 3 C-D). The time WT mice spent with O2 (p<0.0001; t=6,78) and O3 (p<0.0001; t=5,96) was not changed by the EE protocol (Figures 3 C-D).

**Figure 3.**
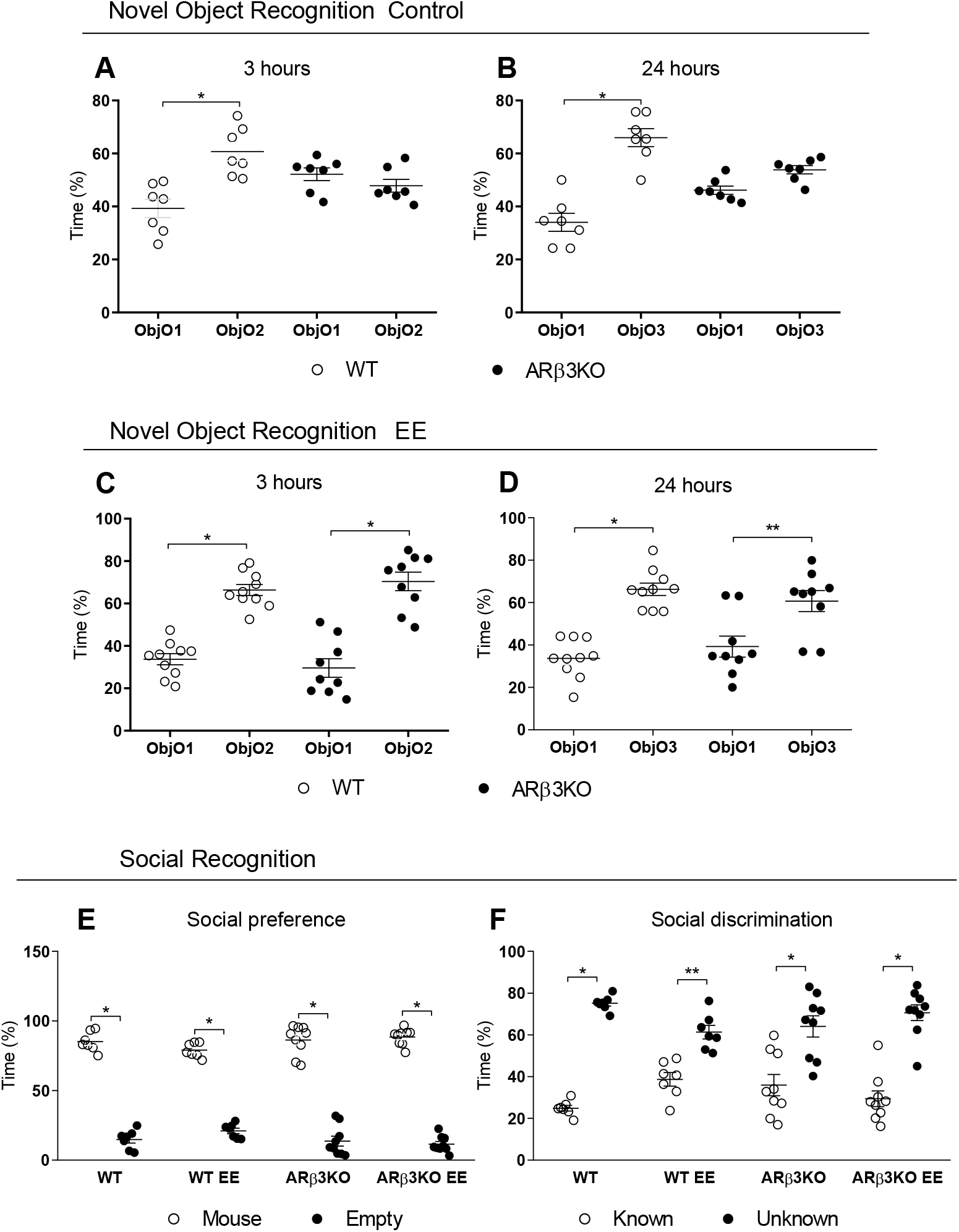
EE exposure early in life corrects cognitive impairment in young adult B3-ARKO mice: *NOR test in mice without exposure to EE*. **(A)** 3 hours after object familiarization, WT mice spent significantly more time with a novel object (O2) than a familiar object (O2), while Adrβ_3_KO mice spent an equal amount of time with both O1 and O2. * p≤0,0001. **(B)** 24 hours after object familiarization, WT mice spent significantly more time with a novel object (O3) than a familiar object O1), while Adr β_3_KO mice spent an equal amount of time with both O1 and O3. * p≤0,0001. *NOR test in mice with exposure to EE late in life*. **(C)** 3 hours after object familiarization both WT and ARB3KO mice exposed to EE early in life spent significantly more time with a novel object (O2) than a familiar object (O1) cognition. * p≤0,0001. **(D)** 24 hours after object familiarization both WT and ARB3KO mice exposed to EE early in life spent significantly more time with a novel object (O3) than a familiar object (O1). * p≤0,0001 and **p=0,019. *SR test*. **(E)** Both WT and ARB3KO mice showed normal preference for social interaction and spent significantly more time in the chamber with a mouse than with a empty cage regardless if they were expose to EE. * p≤0,0001. **(F)** Both WT and ARB3KO mice showed normal preference for social novelty and spent significantly more time in the chamber with a unknown mouse than in the chamber with the now-familiar mouse. * p≤0,0001 and **p=0,001. Values are expressed as mean ± SEM (n=10).

In the SR test, all groups explored the empty cages similarly during the familiarization period (Supp Fig. 1C). The results from the social preference of SR tests showed that WT (p<0.0001; t=17.3) and Adrβ_3_KO (p<0.0001; t=20.28) animals are more interested in spending time with a conspecific animal than with an empty cage (Figure 3E). The exposure to the EE protocol did not change the social preference in both WT EE (p<0.0001; t=14.28) and Adrβ_3_KO (p<0.0001; t=21.5) (Figure 3E). The results obtained from the social discrimination phase of SR test showed that both WT (p<0.0001; t=8.52) and Adrβ_3_KO (p<0.0001; t=5.41) mice not exposed to EE do remember the known animal to whom they were exposed earlier since they spent more time with the unknown mice than with the known mice (Figure 3 F). The exposure to EE protocol did not change this behavior, since WT EE (p=0.001; t=3.84) and Adrβ_3_KO (p<0.0001; t=7.92) spent more time with the unknown mice than with the known one (Figure 3F). The difference in the performance of Adrβ_3_KO mice in NOR compared to SR is explained by the fact that SR test uses conspecific animals, and thus, memory formation is strengthened by stimulus valence.

### EE exposure late in life changes ambulatory and exploratory activity of Adrβ_3_KO and WT mice

Both WT and Adrβ_3_KO adult mice show similar ambulatory activity when exposed to the open field test without effect of time for both groups (F(2,39) = 1.024; p=0.37) (Figure 4A). However, EE increased the ambulatory activity in adult Adrβ_3_KO mice when compared to adult WT mice exposed to EE (F (1,18) = 7.91; p=0.0012) (Figure 4B). EE increased the exploratory activity in WT mice in day 1 (p=0.003), in day 2 (p=0.02), and day 3 (p=0.0003) of testing (Figure 4 C-E). However, EE decreased in Adrβ_3_KO mice in day 1 (p=0.01), in day 2 (p=0.02), and day 3 (p=0.02) of testing (Figure 4C-E). Notably, control Adrβ_3_KO adult mice explored significantly more than control WT adult mice also not exposed to EE in day 2 (p=0.008) and day 3 (p=0.002).

**Figure 4.**
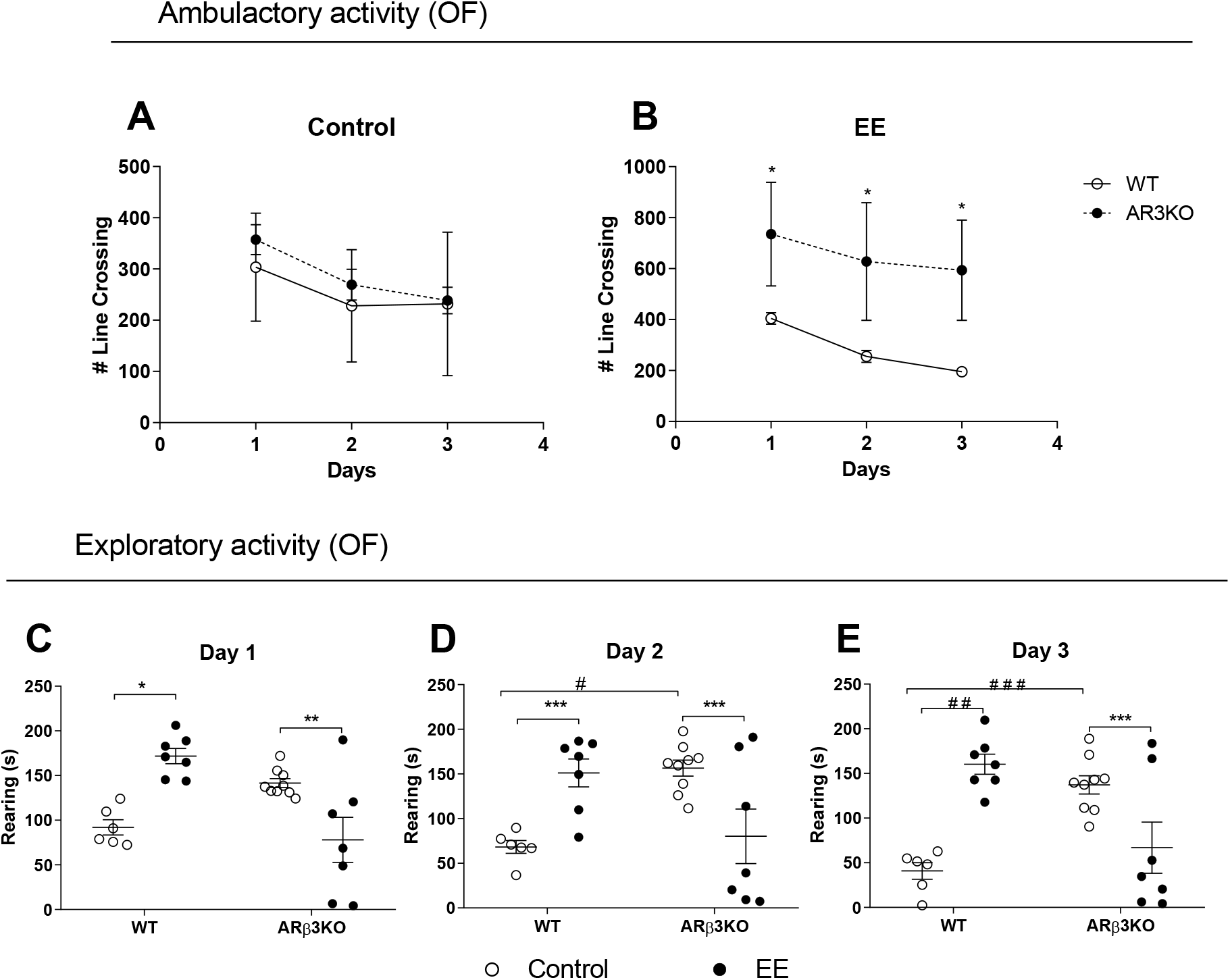
Effect of exposure to EE late in life on locomotor activity and anxiety behavior in young adult B3-ARKO and WT mice. *Open Field*. **(A)** Total number of line crossing of WT and ARB3KO without exposure to EE; **(B)** Total number of line crossing of WT and ARB3KO with exposure to EE; **(C-E)** Total number of rearing. Values are expressed as mean ± SEM (n=10). *p=0.003; **p=0.01; ***p=0.02, #p=0.008, ##p=0.0003 and ###p=0.002.

### EE exposure late in life does not correct cognitive impairment in adults B3-ARKO mice

In the NOR test both groups explored the objects similarly during the familiarization period regardless of the EE exposure and the genotype (Supp. Figure 1D-E). Control and EE exposed WT adult mice spent significantly more time with the new object 3h (O2) (p<0.0001; t=8.88) and 24 h (O3) (p<0.0001; t=6.04) after the familiarization period (Figures 5A-D). In contrast, Control and EE exposed Adrβ_3_KO mice spent similar amounts of time with old (O1) and new objects 3h (O2) (p=0.99; t=0.31) and 24 h (O3) (p=0.21; t=1.68) after the familiarization period (Figure 5A-D). These results show that adult WT mice retained their memory consolidation abilities regardless of older age and show that older Adrβ_3_KO mice exhibited similar memory impairment as younger ones. Remarkably, the exposure of EE later in life at PND120 (Figure 1B) was not able to correct the memory impairment observed in Adrβ_3_KO (Figure 5C-D).

**Figure 5.**
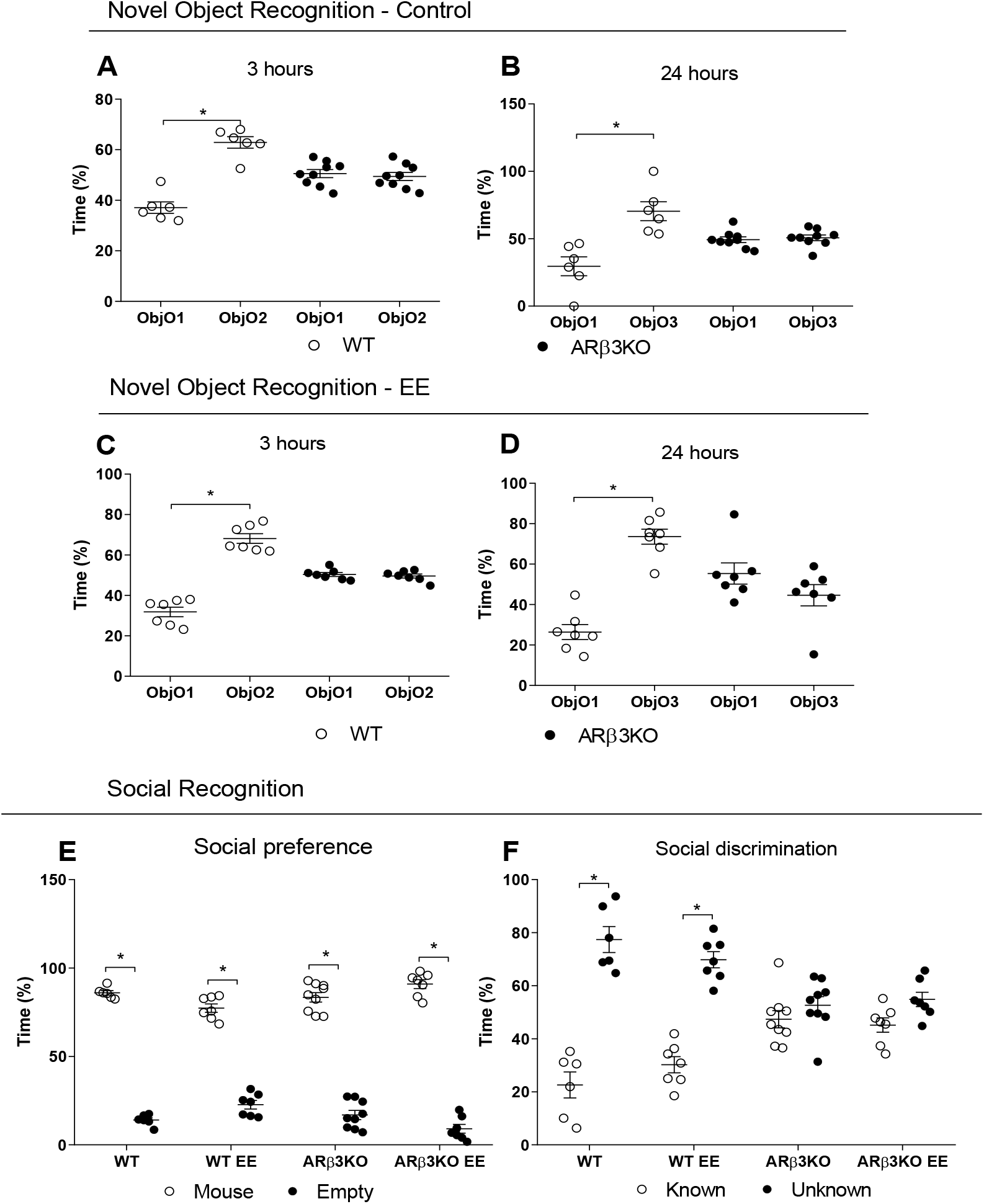
EE exposure late in life does not correct cognitive impairment in adult B3-ARKO mice: *NOR test in mice without exposure to EE*. **(A)** 3 hours after object familiarization, WT mice spent significantly more time with a novel object (O2) than a familiar object (O1), while ARβ_3_KO mice spent an equal amount of time with both O1 and O2. *p<0,0001. **(B)** 24 hours after object familiarization, WT mice spent significantly more time with a novel object (O3) than a familiar object (O1), while ARβ_3_KO mice spent an equal amount of time with both O1 and O3. p≤0,0001*. NOR test in mice with exposure to EE late in life*. **(C)** 3 hours after object familiarization WT spent significantly more time with a novel object (O2) than a familiar object (O1) while ARβ_3_KO mice spent an equal amount of time with both O1 and O2. *p≤0,0001. **(D)** 24 hours after object familiarization WT spent significantly more time with a novel object (O3) than a familiar object (O1) while ARβ_3_KO mice spent an equal amount of time with both O1 and O3. *p≤0,0001. *SR test*. **(E)** Both WT and ARB3KO mice showed normal preference for social interaction and spent significantly more time in the chamber with a mouse than with an empty cage regardless if they were expose to EE. *p≤0,0001. **(F)** WT mice showed normal preference for social novelty and spent significantly more time in the chamber with an unknown mouse than in the chamber with the known mouse, while ARβ_3_KO mice spent an equal amount of time with both known and unknown mice. *p≤0,0001. Values are expressed as mean ± SEM (n=10).

In the SR test, all groups explored the empty cages similarly during the familiarization period (Supp Fig. 1F). The social preference in SR tests showed that control WT and Adrβ_3_KO adult mice spent more time with a conspecific animal than with an empty cage (p<0.0001) (Figure 5E). The control WT adult mice do remember the known animal to whom they were exposed since they spent more time with the unknown mice than with the known one (p<0.0001; t=10.21) and the similar results were observed in EE WT adult mice (p<0.0001; t=7.95) (Figure 5F). However, control Adrβ_3_KO adult mice spent the same amount of time with known and unknown conspecific animal (p=0.94; t=1.2) suggesting a worsen in memory impairment as they grow older, since younger Adrβ_3_KO mice spent more time with the unknown conspecific (Figure 5F). Remarkably, EE was not able to correct this impairment (p=0.23; t=1.94), suggesting that there is a time frame to its efficiency.

## DISCUSSION

The present study showed that the EE protocol when applied to Adrβ_3_KO mice immediately after weaning was able to correct the memory impairment observed in these animals but had no effect when applied to adult animals.

The efficacy of the EE protocol in reversing the short- and long-term memory deficit in young mice due to the lack of Adrβ_3_ could be explained by the findings of several previous studies showing that cognitive enrichment in rats contributes to better learning and the performance of declarative and procedural memory tasks. The increase in the dendritic tree in the third layer of the pyramidal neurons of the temporal lobe, changes in synaptic organization and an increase in the amount of intersections in both the basal and apical dendrites suggests a change in plasticity that involves neural reorganization and an increase in the number of neuronal contacts and the formation of a more complex neuronal network (33–35).

Other studies also show that rats and mice submitted to EE present improvement in learning and memory, as well as better performance in cognitive tasks when compared to rodents kept in standard cages. In addition, they also have increased levels of brain-derived neurotrophic factor (BDNF) and NGF, particularly in the hippocampus region, indicating neuronal growth and proliferation, and brain plasticity (36–38). Thus, it is likely that the improvement in memory tasks observed in young Adrβ_3_KO mice submitted to EE can be explained by these changes in neurotrophin expression.

The constant exposure of animals to novelty through EE involves some degree of stress in animals. The animals were exposed to various stimuli, including those of negative valence, such as bedding with the smell of rats. When the animal has contact with stimuli with negative valence, it could induce an increase in the peripheral release of adrenaline that activates the expression of Adrβs in the vagus nerve, which will in turn activate the locus coeruleus (LC). The activation of the LC leads to an increase in NE release in the hippocampus and amygdala (39, 40). Thus, it is possible that the increase in NE levels induced by EE activates a compensatory pathway through β-ARs, reversing the memory deficit in young Adrβ_3_KO mice.

Although EE was very beneficial for memory when applied to young animals, the results obtained when EE was introduced to older Adrβ_3_KO mice showed that the protocol used is not able to reverse the damage caused by the absence of ARβ_3_ when it starts in adulthood. In fact, aging worsens the memory of Adrβ_3_KO mice, since their good performance in the social recognition test at 2-3 months of age is not repeated at 6-7 months of age, despite the amygdala activation due to the valence of the stimulus. It has been shown that LC degeneration is a common neuropathological feature of neurogenerative diseases such as Alzheimer’s (41, 42). In fact, early degeneration of the LC could trigger or be involved in the progression of neurogenerative diseases (43). The fact that a lack of Adrβ_3_KO leads to a greater loss in cognition highlights the role of the noradrenergic signaling pathway in the course of dementia. Also, it has been shown that EE in older healthy mice is not as efficient in improving cognition as it is in younger animals (44).

In healthy rodents, locus coeruleus projections to different brain regions begin to decline by 7–15 months of age (45, 46). Other studies with rodents and primates have found a correlation between memory loss and the increasing appearance of lesions, and consequent cell loss in the hippocampus and entorhinal cortex, with age (47, 48). Advancing age leads to a loss of 10 to 20% of brain mass when compared to a young brain. This can lead to variations in cell loss in different brain regions and, consequently, more serious losses in certain regions than in others (49). The lack of Adrβ_3_ combined with the functional changes typical of advancing age can aggravate damage to memory formation processes and that could be the mechanism underling the worsen in memory observed in adult Adrβ_3_KO mice. The effects of aging may explain the absence of any benefit from EE when applied to older animals. The results of the present study reinforce the idea that early stimulation of individuals is beneficial for cognition and can prevent or delay early memory impairment caused by defects in neuronal signaling involved in cognition.

EE does not alter locomotor capacity when applied to young animals regardless of the genotype but increased ambulatory activity in older Adrβ_3_KO mice. This suggests that the stimulus represented by the EE may improve the activity of animals at an older age. The influence of EE on the exploratory behavior of mice has already been evaluated in other studies, but there is as yet no consensus on its influence (50, 51).

In conclusion, the results obtained show that Adrβ_3_ has an important role in memory as aging leads to a worsening in the memory of Adrβ_3_KO animals that is not corrected by the EE protocol used in this study. However, the EE protocol can reverse memory damage in young Adrβ_3_KO animals. Further studies are needed to understand the role of Adrβ_3_ in aging, and the mechanism involved in the effect of EE in reversing memory impairment.

## Acknowledgement

Support from FAPESP 2017/18277-0 for MOR; PROEX 1133/2019 for MOR; CAPES for TTR; FAPESP 2017/04491-0 for BPPN.

## REFERENCES

1. McGaugh JL. The amygdala modulates the consolidation of memories of emotionally arousing experiences. Annu Rev Neurosci. 2004;27:1–28.

2. Lu B, Pang PT, Woo NH. The yin and yang of neurotrophin action. Nat Rev Neurosci. 2005;6(8):603–14.

3. Hillman KL, Knudson CA, Carr PA, Doze VA, Porter JE. Adrenergic receptor characterization of CA1 hippocampal neurons using real time single cell RT-PCR. Brain Res Mol Brain Res. 2005;139(2):267–76.

4. Hillman KL, Doze VA, Porter JE. Functional characterization of the beta-adrenergic receptor subtypes expressed by CA1 pyramidal cells in the rat hippocampus. J Pharmacol Exp Ther. 2005;314(2):561–7.

5. Cox DJ, Racca C, LeBeau FE. Beta-adrenergic receptors are differentially expressed in distinct interneuron subtypes in the rat hippocampus. J Comp Neurol. 2008;509(6):551–65.

6. O’Dell TJ, Connor SA, Guglietta R, Nguyen PV. beta-Adrenergic receptor signaling and modulation of long-term potentiation in the mammalian hippocampus. Learn Mem. 2015;22(9):461–71.

7. Izquierdo I, Bevilaqua LR, Rossato JI, Bonini JS, Medina JH, Cammarota M. Different molecular cascades in different sites of the brain control memory consolidation. Trends Neurosci. 2006;29(9):496–505.

8. Cammarota M, Bevilaqua LR, Rossato JI, Lima RH, Medina JH, Izquierdo I. Parallel memory processing by the CA1 region of the dorsal hippocampus and the basolateral amygdala. Proc Natl Acad Sci U S A. 2008;105(30):10279–84.

9. Kemp A, Manahan-Vaughan D. Beta-adrenoreceptors comprise a critical element in learning-facilitated long-term plasticity. Cereb Cortex. 2008;18(6):1326–34.

10. Straube T, Korz V, Balschun D, Frey JU. Requirement of beta-adrenergic receptor activation and protein synthesis for LTP-reinforcement by novelty in rat dentate gyrus. J Physiol. 2003;552(Pt 3):953–60.

11. Hagena H, Manahan-Vaughan D. Learning-facilitated long-term depression and longterm potentiation at mossy fiber-CA3 synapses requires activation of beta-adrenergic receptors. Frontiers in integrative neuroscience. 2012;6:23.

12. Hansen N, Manahan-Vaughan D. Hippocampal long-term potentiation that is elicited by perforant path stimulation or that occurs in conjunction with spatial learning is tightly controlled by beta-adrenoreceptors and the locus coeruleus. Hippocampus. 2015;25(11):1285–98.

13. Souza-Braga P, Lorena FB, Nascimento BPP, Marcelino CP, Ravache TT, Ricci E, et al. Adrenergic receptor beta3 is involved in the memory consolidation process in mice. Braz J Med Biol Res. 2018;51(10):e7564.

14. Mellor D, Hunt S, Gusset M. Caring for Wildlife: The World Zoo and Aquarium Animal Welfare Strategy 2015.

15. Bondi CO, Klitsch KC, Leary JB, Kline AE. Environmental enrichment as a viable neurorehabilitation strategy for experimental traumatic brain injury. Journal of neurotrauma. 2014;31(10):873–88.

16. Segovia G, del Arco A, Mora F. Environmental enrichment, prefrontal cortex, stress, and aging of the brain. J Neural Transm (Vienna). 2009;116(8):1007–16.

17. Leggio MG, Mandolesi L, Federico F, Spirito F, Ricci B, Gelfo F, et al. Environmental enrichment promotes improved spatial abilities and enhanced dendritic growth in the rat. Behavioural brain research. 2005;163(1):78–90.

18. Zimmermann A, Stauffacher M, Langhans W, Wurbel H. Enrichment-dependent differences in novelty exploration in rats can be explained by habituation. Behavioural brain research. 2001;121(1-2):11–20.

19. Gonzalez-Pardo H, Arias JL, Vallejo G, Conejo NM. Environmental enrichment effects after early stress on behavior and functional brain networks in adult rats. PloS one. 2019;14(12):e0226377.

20. Dandi E, Kalamari A, Touloumi O, Lagoudaki R, Nousiopoulou E, Simeonidou C, et al. Beneficial effects of environmental enrichment on behavior, stress reactivity and synaptophysin/BDNF expression in hippocampus following early life stress. Int J Dev Neurosci. 2018;67:19–32.

21. von Bohlen Und Halbach O, von Bohlen Und Halbach V. BDNF effects on dendritic spine morphology and hippocampal function. Cell Tissue Res. 2018;373(3):729–41.

22. Robison LS, Francis N, Popescu DL, Anderson ME, Hatfield J, Xu F, et al. Environmental Enrichment: Disentangling the Influence of Novelty, Social, and Physical Activity on Cerebral Amyloid Angiopathy in a Transgenic Mouse Model. International journal of molecular sciences. 2020;21(3).

23. Wei Z, Meng X, El Fatimy R, Sun B, Mai D, Zhang J, et al. Environmental enrichment prevents Abeta oligomer-induced synaptic dysfunction through mirna-132 and hdac3 signaling pathways. Neurobiol Dis. 2020;134:104617.

24. Susulic VS, Frederich RC, Lawitts J, Tozzo E, Kahn BB, Harper ME, et al. Targeted disruption of the beta 3-adrenergic receptor gene. J Biol Chem. 1995;270(49):29483–92.

25. Simpson J, Kelly JP. The impact of environmental enrichment in laboratory rats--behavioural and neurochemical aspects. Behavioural brain research. 2011;222(1):246–64.

26. Huttenrauch M, Salinas G, Wirths O. Effects of Long-Term Environmental Enrichment on Anxiety, Memory, Hippocampal Plasticity and Overall Brain Gene Expression in C57BL6 Mice. Frontiers in molecular neuroscience. 2016;9:62.

27. Seibenhener ML, Wooten MC. Use of the Open Field Maze to measure locomotor and anxiety-like behavior in mice. J Vis Exp. 2015(96):e52434.

28. Hall C, Ballachey, E.L.. A study of the rat’s behavior in a field. A contribution to method in comparative psychology. University of California Publications in psychology. 1932;6:1–12.

29. Leger M, Quiedeville A, Bouet V, Haelewyn B, Boulouard M, Schumann-Bard P, et al. Object recognition test in mice. Nature protocols. 2013;8(12):2531–7.

30. Moy SS, Nadler JJ, Young NB, Nonneman RJ, Segall SK, Andrade GM, et al. Social approach and repetitive behavior in eleven inbred mouse strains. Behavioural brain research. 2008;191(1):118–29.

31. Crawley JN. Mouse behavioral assays relevant to the symptoms of autism. Brain pathology. 2007;17(4):448–59.

32. Novaes GF, Amado D, Scorza FA, Cysneiros RM. Social behavior impairment in offspring exposed to maternal seizures in utero. J Neural Transm (Vienna). 2012;119(6):639–44.

33. Moreno-Jimenez EP, Jurado-Arjona J, Avila J, Llorens-Martin M. The Social Component of Environmental Enrichment Is a Pro-neurogenic Stimulus in Adult c57BL6 Female Mice. Front Cell Dev Biol. 2019;7:62.

34. Kempermann G. Environmental enrichment, new neurons and the neurobiology of individuality. Nat Rev Neurosci. 2019;20(4):235–45.

35. Duran-Carabali LE, Arcego DM, Sanches EF, Odorcyk FK, Marques MR, Tosta A, et al. Preventive and therapeutic effects of environmental enrichment in Wistar rats submitted to neonatal hypoxia-ischemia. Behavioural brain research. 2019;359:485–97.

36. Barros W, David M, Souza A, Silva M, Matos R. Can the effects of environmental enrichment modulate BDNF expression in hippocampal plasticity? A systematic review of animal studies. Synapse. 2019;73(8):e22103.

37. Zhang XQ, Mu JW, Wang HB, Jolkkonen J, Liu TT, Xiao T, et al. Increased protein expression levels of pCREB, BDNF and SDF-1/CXCR4 in the hippocampus may be associated with enhanced neurogenesis induced by environmental enrichment. Molecular medicine reports. 2016;14(3):2231–7.

38. Leger M, Paizanis E, Dzahini K, Quiedeville A, Bouet V, Cassel JC, et al. Environmental Enrichment Duration Differentially Affects Behavior and Neuroplasticity in Adult Mice. Cereb Cortex. 2015;25(11):4048–61.

39. Miyashita T, Williams CL. Peripheral arousal-related hormones modulate norepinephrine release in the hippocampus via influences on brainstem nuclei. Behavioural brain research. 2004;153(1):87–95.

40. Jankord R, Herman JP. Limbic regulation of hypothalamo-pituitary-adrenocortical function during acute and chronic stress. Ann N Y Acad Sci. 2008;1148:64–73.

41. McMillan PJ, White SS, Franklin A, Greenup JL, Leverenz JB, Raskind MA, et al. Differential response of the central noradrenergic nervous system to the loss of locus coeruleus neurons in Parkinson’s disease and Alzheimer’s disease. Brain Res. 2011;1373:240–52.

42. Weinshenker D. Long Road to Ruin: Noradrenergic Dysfunction in Neurodegenerative Disease. Trends Neurosci. 2018;41(4):211–23.

43. Pamphlett R. Uptake of environmental toxicants by the locus ceruleus: a potential trigger for neurodegenerative, demyelinating and psychiatric disorders. Medical hypotheses. 2014;82(1):97–104.

44. Chandler K, Dosso H, Simard S, Siddiqi S, Rudyk C, Salmaso N. Differential Effects of Short-term Environmental Enrichment in Juvenile and Adult Mice. Neuroscience. 2020;429:23–32.

45. Shirokawa T, Ishida Y, Isobe KI. Age-dependent changes in axonal branching of single locus coeruleus neurons projecting to two different terminal fields. Journal of neurophysiology. 2000;84(2):1120–2.

46. Ishida Y, Shirokawa T, Miyaishi O, Komatsu Y, Isobe K. Age-dependent changes in projections from locus coeruleus to hippocampus dentate gyrus and frontal cortex. European Journal of Neuroscience. 2000;12(4):1263–70.

47. Haberman RP, Colantuoni C, Stocker AM, Schmidt AC, Pedersen JT, Gallagher M. Prominent hippocampal CA3 gene expression profile in neurocognitive aging. Neurobiol Aging. 2011;32(9):1678–92.

48. Almaguer-Melian W, Cruz-Aguado R, Riva Cde L, Kendrick KM, Frey JU, Bergado J. Effect of LTP-reinforcing paradigms on neurotransmitter release in the dentate gyrus of young and aged rats. Biochem Biophys Res Commun. 2005;327(3):877–83.

49. Gannon M, Che P, Chen Y, Jiao K, Roberson ED, Wang Q. Noradrenergic dysfunction in Alzheimer’s disease. Frontiers in neuroscience. 2015;9:220.

50. Kazlauckas V, Pagnussat N, Mioranzza S, Kalinine E, Nunes F, Pettenuzzo L, et al. Enriched environment effects on behavior, memory and BDNF in low and high exploratory mice. Physiol Behav. 2011;102(5):475–80.

51. Amaral OB, Vargas RS, Hansel G, Izquierdo I, Souza DO. Duration of environmental enrichment influences the magnitude and persistence of its behavioral effects on mice. Physiol Behav. 2008;93(1-2):388–94.

52. Gong X, Chen Y, Chang J, Huang Y, Cai M, Zhang M. Environmental enrichment reduces adolescent anxiety- and depression-like behaviors of rats subjected to infant nerve injury. Journal of neuroinflammation. 2018;15(1):262.

53. Novaes LS, Dos Santos NB, Batalhote RFP, Malta MB, Camarini R, Scavone C, et al. Environmental enrichment protects against stress-induced anxiety: Role of glucocorticoid receptor, ERK, and CREB signaling in the basolateral amygdala. Neuropharmacology. 2017;113(Pt A):457–66.

54. Mora-Gallegos A, Fornaguera J. The effects of environmental enrichment and social isolation and their reversion on anxiety and fear conditioning. Behavioural processes. 2019;158:59–69.

55. Chaudhuri A, Zangenehpour S, Rahbar-Dehgan F, Ye F. Molecular maps of neural activity and quiescence. Acta neurobiologiae experimentalis. 2000;60(3):403–10.

56. Rao RP, Anilkumar S, McEwen BS, Chattarji S. Glucocorticoids protect against the delayed behavioral and cellular effects of acute stress on the amygdala. Biol Psychiatry. 2012;72(6):466–75.

