## Supplementary figures and images for "Environmental enrichment reverses memory impairment in B3-ARKO mice"

### Supplemental Figure 1

#### Slide 1
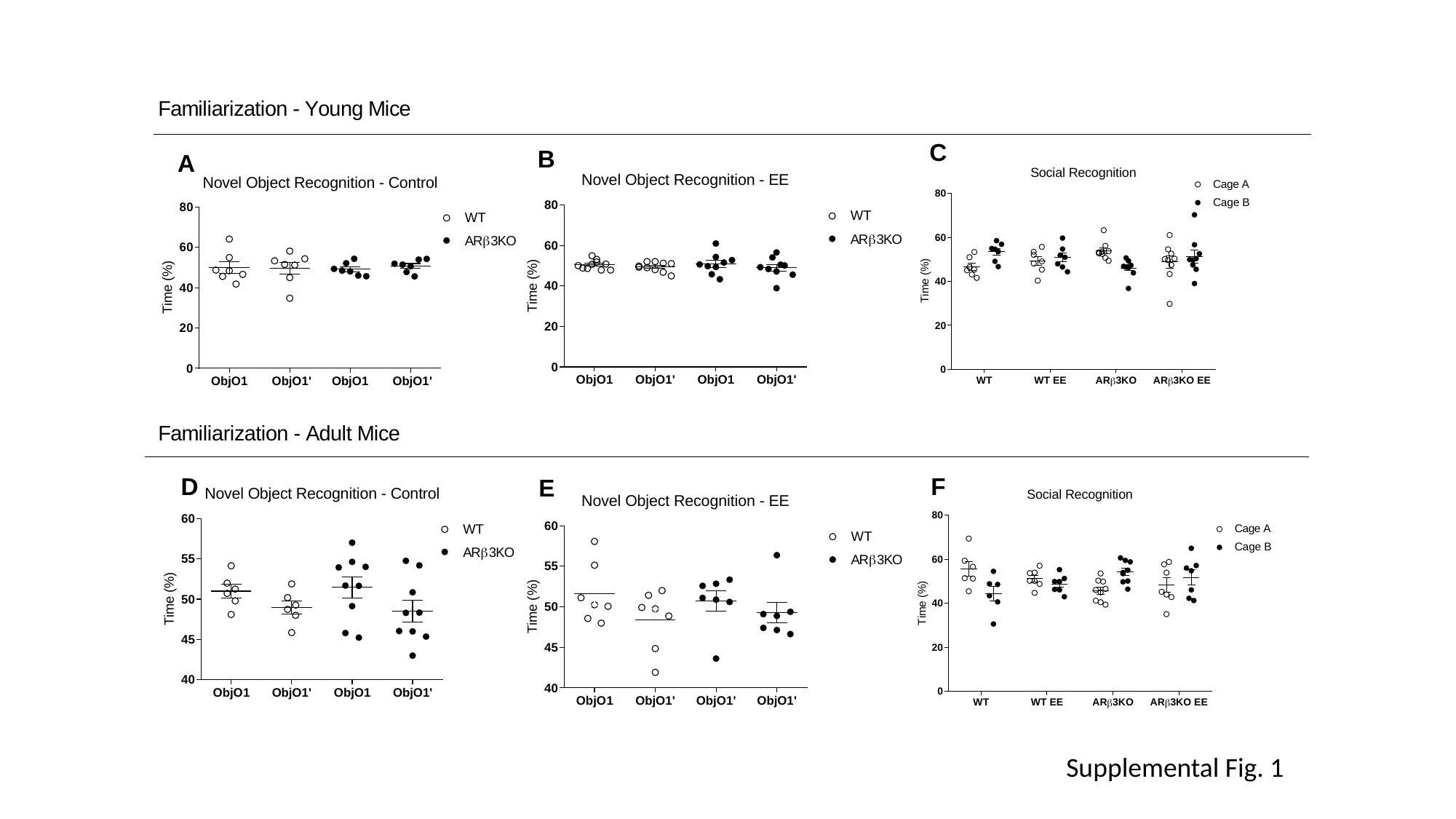

Supplemental Fig. 1
